# Bessel beam tomography for fast volume Imaging

**DOI:** 10.1101/552661

**Authors:** Andres Flores Valle, Johannes D. Seelig

## Abstract

Light microscopy on dynamic samples, for example neural activity in the brain, requires imaging large volumes at high rates. Here, we develop a tomography approach for scanning fluorescence microscopy which allows recording volume images at frame scan rates. Volumes are imaged by simultaneously recording four independent projections at different angles using temporally multiplexed, tilted Bessel beams. From the resulting projections, volumes are reconstructed using inverse Radon transforms combined with three dimensional convolutional neural networks (U-net). This tomography approach is suitable for experiments requiring fast volume imaging of sparse samples, as for example often encountered when imaging neural activity in the brain.

## 1. Introduction

In many fluorescence microscopy experiments samples show dynamics which evolves not in a single focal plane but in a volume. For example, imaging neural activity in the brain requires monitoring multiple cells that are distributed in three dimensions [1, 2]. In this situation, the temporal resolution of the microscope needs to match the time scale of the sample dynamics, which for neural activity monitored with calcium indicators, lies in the range of several tens of Hertz [1, 3, 4].

While such imaging rates are standard for scanning a single plane, they are more difficult to reach for volumes. The scan rate is ultimately limited by the requirement of integrating a sufficient number of photons per scanned pixel. To overcome this limit, solutions for volume imaging have been developed that scan multiple pixels in parallel [3, 4]. Multifocal microscopy, for example, images several spots at the same time, but requires detectors with multiple pixels making this approach susceptible to scattering (for example when imaging in tissue) [3, 5]. Multiple focal spots can also be imaged in parallel with single pixel detectors: by introducing temporal offsets between different (typically pulsed) beams, fluorescence emission can be sorted into different temporal channels [6, 7].

To speed up volume imaging rates, extended volumes - instead of multiple focal spots - can be excited at the same time, for example using Bessel beams, line foci or light sheets. Combined with camera detectors, these approaches work best for weakly scattering samples [3, 4], but Bessel beams or line foci [5] can also be used for imaging with non-descanned PMT detection. This is more suitable for imaging in scattering tissue and results in a projection of the entire imaging volume along the beam axis [8-12]. While for Bessel beams this approach offers the advantage of recording from an entire volume in a single frame scan, it comes at the cost of loosing all axial resolution.

Axial resolution was restored in experiments that used two Bessel beams which imaged the sample at different angles [9, 10, 12] for generating two different projections. These projections were either recorded sequentially [10] or simultaneously (as required for moving samples such as brain tissue *in vivo)* in a single channel by averaging over two different projections [12]. From these projections axial information was recovered based on the distance between prominent, sparse features such as fluorescent beads or cell bodies [10, 12].

Recording multiple projections at different viewing angles generally forms the basis for reconstructing volumes in tomographic imaging [13]. While such tomographic approaches have been implemented for fluorescence microscopy by rotating the samples with respect to the illumination direction [14, 15], this is not compatible with most *in vivo* imaging preparations. A tomography approach compatible with fluorescence microscopy for *in vivo* imaging has recently been developed with Gaussian line foci generating projections in the focal plane and was combined with scanning the objective along the axial direction to record volumes [5].

We here develop a tomography method compatible with an *in vivo* imaging configuration based on Bessel beams. Volume information is obtained from four independent projections recorded from four different angles using temporally multiplexed, tilted Bessel beams in a single frame scan. For volume reconstruction we combine inverse Radon transforms adapted for Bessel beam scanning with machine learning [13, 16-20]. Machine learning has been shown for example in optical phase imaging to allow high resolution reconstruction from sparse projections at shallow angles similar to the ones used here [18]. We show that three-dimensional convolutional neural networks (U-net) trained on suitable datasets similarly improve reconstructions in Bessel beam tomography.

The method presented here allows volumetric imaging with non-descanned detection in a single frame scan which makes it suitable for experiments that require imaging of sparse samples at fast volume rates, such as neural activity from cell bodies in the brain.

## 2. Results

### 2.1. Setup

Four different projections were generated by using four different Bessel beams tilted with an angle of 10 degrees with respect to the optical axis and an azimuth separation of 90 degrees between neighboring beams. The setup is shown in Fig. 1 a. A laser beam (Coherent Discovery, 120 fs pulse width) is split into four beams with non-polarizing plate beam splitters and the (nevertheless polarization dependent) splitting ratio is additionally balanced using a half-wave plate for each beam (not shown). The beams are separated with a delay of 3 ns (corresponding to a 1 m additional path length per beam) [6, 7] which allows distinguishing the different projections with a single detector using photon counting at a time resolution faster than the pulse repetition rate, provided that the fluorescence lifetime is shorter than the pulse separation [6]. The beam width and collimation is adjusted for each beam with a telescope (two achromatic lenses, one mounted on a translation stage).

**Fig. 1.**
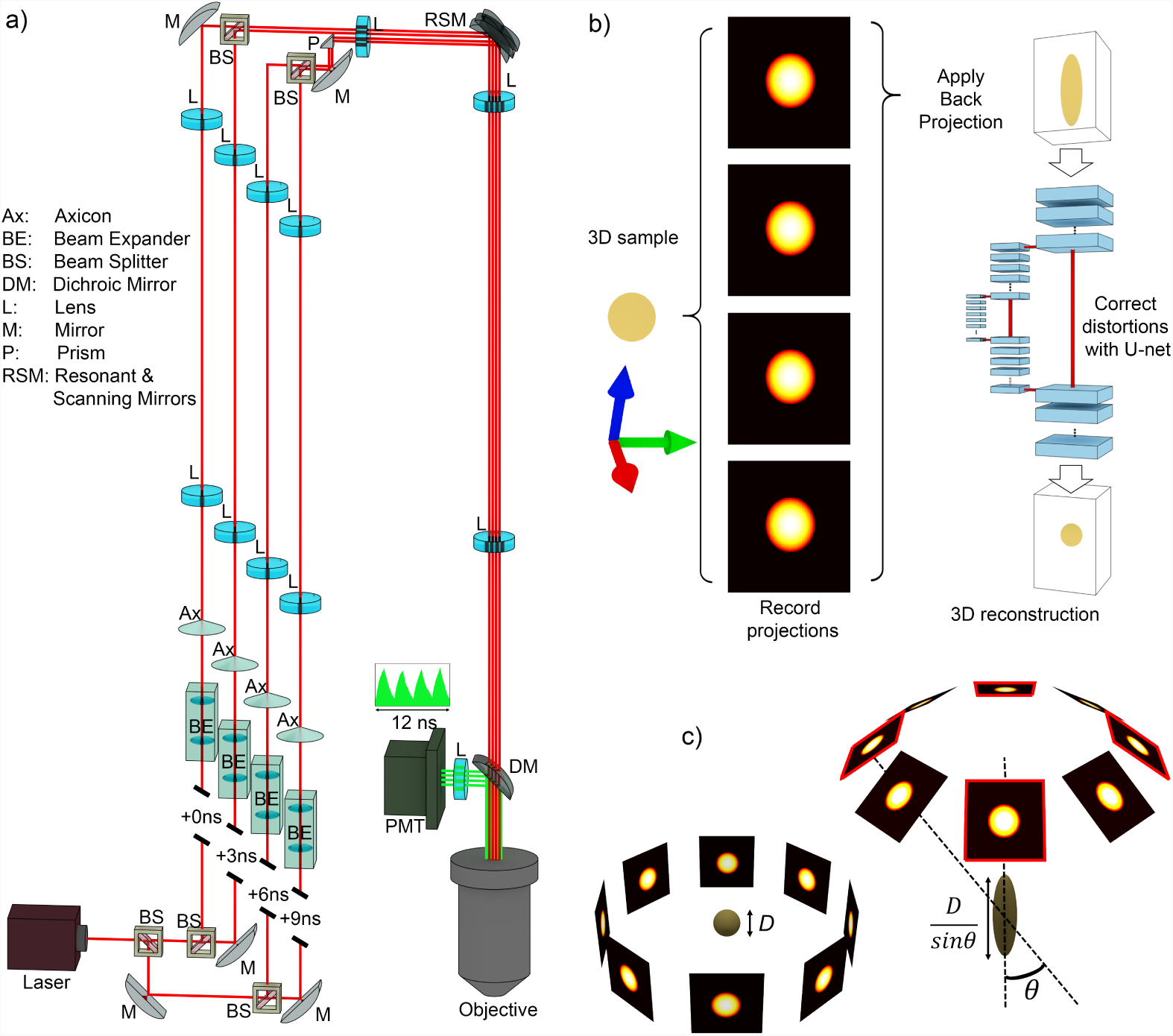
**a)** Schematic of the experimental setup. A laser beam is split into four temporally offset beams using beam splitters (BS). Each beam is converted into a Bessel beam with independently adjusted beam parameters. The Bessel beams excite the volume from four different sides. Inset: Fluorescence emission is detected in four temporally separated channels. **b)** Workflow for volumetric reconstruction: four projections are recorded from a three dimensional (3D) sample and back projection is applied to obtain its 3D tomographic reconstruction. Then, a U-net is applied to correct distortions in the reconstructed sample. **c)** Illustration of 3D tomography with different viewing angles. Left side: projections parallel to the *xy* plane allow a perfect reconstruction of a sphere of diameter *D*. Right side: projections at an oblique angle *θ* produce a reconstructed sphere elongated in the *z* axis by a factor of 1/sin(*θ*) even when using more than four (red frames) projections. This is corrected for by using convolutional networks.

Each beam is sent through an axicon (AX252-B, all optical components were from Thorlabs unless otherwise noted) for generating a Bessel beam. The beam is collimated after the axicon with an achromatic lens (AC254-100-B) and the resulting ring is imaged onto the back focal plane of the microscope objective. For this, the ring is first demagnified and imaged onto the scan mirrors using a telescope (AC254-200-B and AC254-100-B), with the latter lens common to all four beam paths. The scan mirror is imaged onto the back focal plane of the objective using a scan lens (AC300-050-B) and tube lens (AC508-300-B).

To achieve tilted illumination from four opposing azimuth angles (see Fig. 3), the last lens in each beam path (before the beams are recombined) is displaced laterally and/or vertically for each beam, resulting in an off-center Bessel ring in each quadrant of the back focal plane of the objective. The Bessel ring diameters at the back aperture of the objective were slightly below half the objective aperture diameter and allowed laterally displacing the beam without clipping.

Depending on the parameters and methods used to generate Bessel beams, different profiles in terms of width, length and axial intensity distribution result [21]. The axial intensity profile additionally depends on the placement of the collimating lenses [12, 21]. Here, we aimed for a beam length of about 80 *μ*m as a compromise between recording a sufficiently large volume while limiting the power required for two-photon excitation in each beam. The focal length of each beam was iteratively adjusted based on the two-photon point spread function by changing the beam collimation using the translation stage in the beam expander [12].

The four beams are recombined by first combining pairs of beams - after adjusting their polarizations othogonally with a half-wave plate (not shown) - with a polarizing beam splitter cube. All four beams are then combined with a right angle reflecting prism positioned just before the common telescope lens (that forms the demagnified image of the Bessel ring on the scan mirror). The reflecting prism is positioned laterally into the beam path using a translation stage, reflecting two beams at 90 degrees while allowing the other two beams to pass near the edge of the prism. This is possible without clipping any of the beams by laterally displacing the two passing beams in the appropriate neighboring quadrants in the objective back focal plane (as required for tilting of the Bessel beams). An additional Gaussian beam path (not shown) was used for recording reference image stacks. A flip mirror was used to direct the beam through a telescope towards a polarizing beam splitter cube mounted on a magnetic mount that was manually inserted before the resonant scanner.

All experiments were performed with a custom-built two-photon microscope controlled with Scanimage (Vidrio) [22] with a resonant scanner (resonant scan box, Sutter Instruments) and components similar to the ones described in [23]. Scanimage allows FPGA-based (National Instruments, PXIe-7975R) photon counting, in our setup at a time resolution of 1.28 GHz (16 channels between laser pulses at 80 MHz repetition rate).

### 2.2. Tomographic volume reconstruction

A typical 3D tomography approach, for example in medical imaging, acquires 2D projections of a sample by varying the azimuth angle at a fixed elevation of 90 degrees, as illustrated on the left side of Fig. 1 c. Changing the elevation to an oblique angle as shown on the right side of Fig. 1 c distorts the reconstruction by a factor of 1/sin(*θ*). Using tilted Bessel beams to record projections results in a distortion along the *z* axis and this was (partially) corrected for using machine learning. The workflow of our tomography approach is shown in Fig. 1 b: first, four projections are recorded from a 3D sample, second, back projection is applied resulting in a distorted 3D reconstruction and, third, a convolutional network (3D U-net [24, 25]) is used to recover the 3D object. These steps are described in more detail below.

#### 2.2.1. Radon transform for Bessel beam scanning

For volume reconstruction we developed a 2D Radon transform based approach for Bessel beam scanning with tilted beams. In this approach, the amount of light emitted following excitation at a given point **x** ∈ ℝ^3^ in a volumetric sample is described with a function *f* : ℝ^3^ → ℝ. The Bessel beam is approximated as a line segment of length *ℓ* with constant intensity centered at a point **x**_*c*_ ∈ ℝ^3^ and a direction given by the normalized vector **ŝ** ∈ ℝ^3^ with a parameter *t*. The total amount of light emitted by the sample due to excitation by the beam is then described in terms of the Radon transform along the segment:

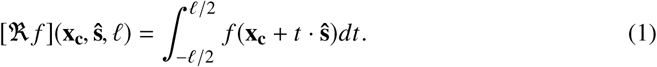

We define an image of size 2*N* × 2*N* as a function *I*: {ℤ ⋂ [−*N, N*]} × {ℤ ⋂ [−*N, N*]} → ℝ representing the intensity of a pixel at a position with integer indices (*v, μ*). To create an image, the Bessel beam is scanned with the two scanning mirrors across the field of view and for each pixel the resulting intensity is recorded. When scanning the tilted Bessel beams we observed slight changes of both the vector **x**_**c**_ and the direction vector **ŝ** across large fields of view compared to a Gaussian reference stack, see Fig. 3. Both vectors are therefore described as vector fields depending on the pixel coordinates (*v, μ*): the offset vector field, **x**_**c**_ = **x**_**c**_(*v, μ, z*), and the the direction vector field, **ŝ** = **ŝ** (*v, μ, z*). A general expression for the intensity in each pixel is then given by:

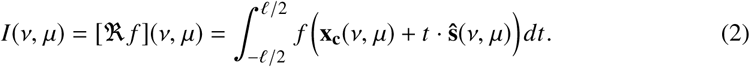

#### 2.2.2. Calibration of Bessel beam projections

The vector fields 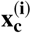 and **ŝ**^**(i)**^ are required for reconstructing the volumetric sample *f* (**x**) from its projections *I*^(*i*)^(*v, μ*; *z*) and are measured experimentally for each projection *I*^(*i*)^ with a volumetric sample of 1 *μ*m diameter fluorescent beads suspended in Agarose (see Fig. 2 a and Fig. 3). For this, a volume *V*^(*i*)^ was recorded in a z-stack while scanning the four Bessel beams resulting in volumes of 2D projections, *V*^(*i*)^(*v, μ, z*) ≡ {*I*^(*i*)^(*v, μ*; *z*)/*z* ∈ [*z*_*min*_, *z*_*max*_]}. The variation of the vector field in z-direction was negligible.

**Fig. 2.**
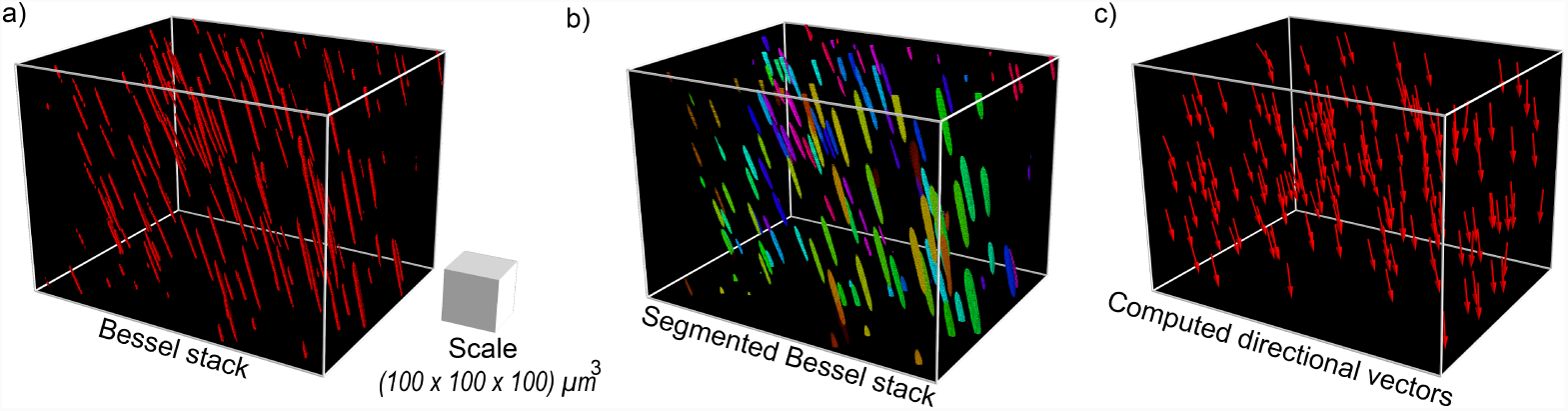
Calibration procedure for a single Bessel beam. **a)** A volumetric sample of 1 μ m diameter beads in Agarose recorded with 1 *μ* m step size in the axial direction (z-axis). **b)** Segmentation of beads to calculate the direction and center of the Bessel beam using Principal Component Analysis (PCA) (segmented beams are colored randomly for visualization). **c)** Computed directional vectors at the center of the beads. These vectors are used in linear regression to compute the directional vector field of the Bessel beam.

The recorded volumes were segmented using a threshold after Gaussian filtering for noise reduction, see Fig. 2 a and b. The center position of each segmented bead 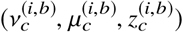 was identified using the *skimage* function *regionprops* and Principal Component Analysis (PCA) was applied to obtain the axis with the highest eigenvalue as the direction vector **ŝ** ^(*i,b*)^ at the center point 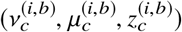 as well as the length of the beam *ℓ*^(*i,b*)^, see Fig. 2 c and Fig. 3.

**Fig. 3.**
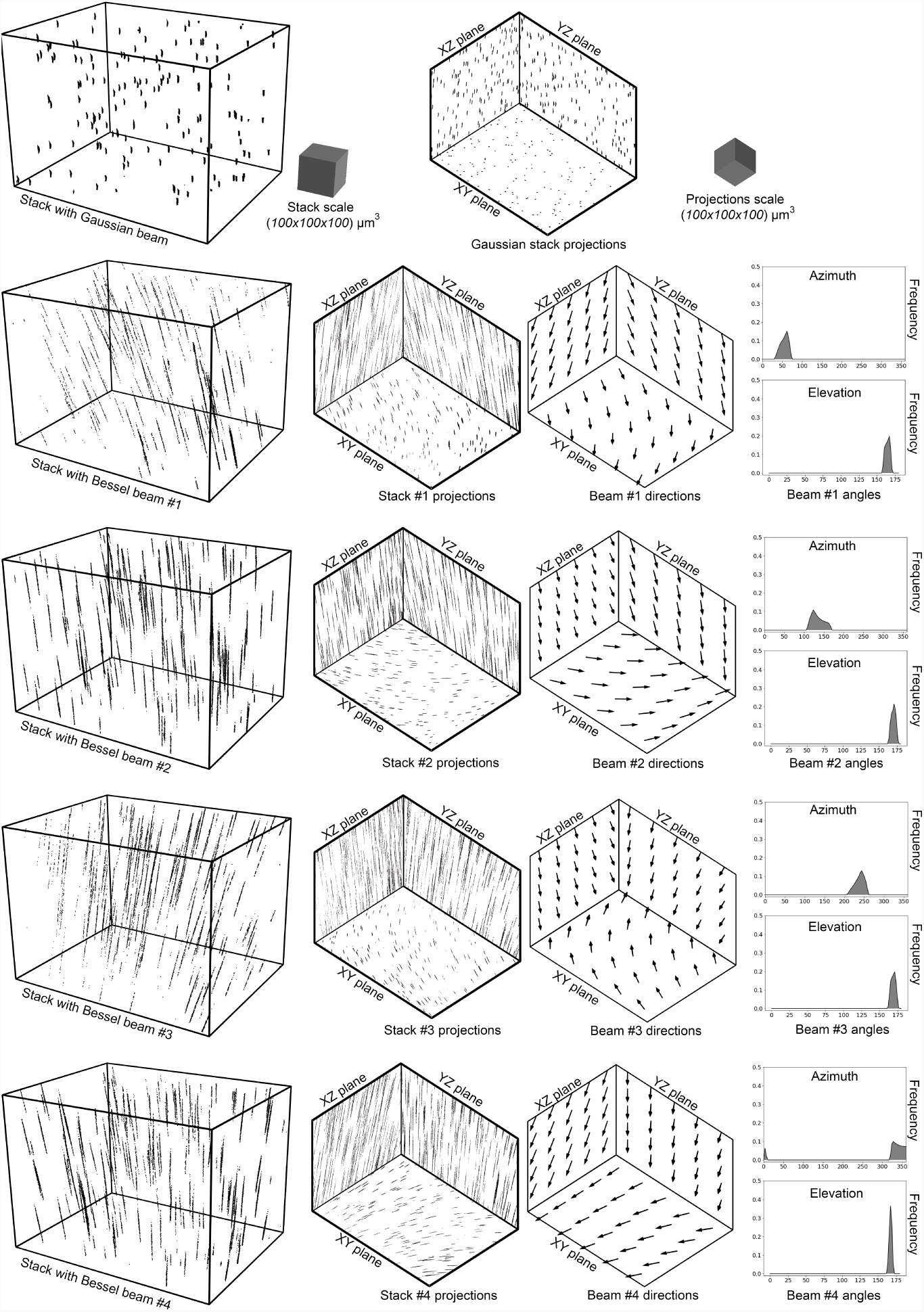
Directional vector fields of each Bessel beam measured and computed from a volumetric sample of 1 *μm* diameter beads. Top row: sample recorded with a Gaussian beam (left side) and projections of the volume in the *xy, xz* and *yz* planes, respectively (right side). Second row to bottom row: volumetric samples recorded by the four Bessel beams (left side) and plane projections (left center). The extrapolated vector field for each beam (right center) is shown in plane projections. The distribution of the elevation and azimuth angles for each directional vector field is shown in the rightmost column. The elevation angle is centered around 170 degrees for all Bessel beams (following the convention of spherical coordinates, corresponding to a 10 degree tilt with respect to the optical axis), while the azimuth separation is approximately 90 degrees between adjacent beams.

Making use of all beads measured in each stack *i* and the corresponding directional vectors,

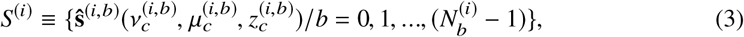

we used linear regression to obtain the directional vector field across the entire field of view (*v, μ, z*,)

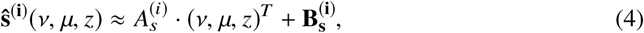

where *A*_*s*_ is a 3 × 3 matrix while **B**_s_ is a bias vector, which are both parameters fit to the data *S*^(*i*)^

The length of the beam in each stack *i* is considered constant, and it is computed as the average value of the lengths obtained for all beads:

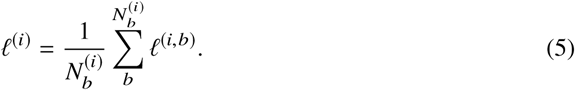

Due to the selected beam length and collimation of the tilted Bessel beams, the different beams don’t meet at the center of the field of view (similar to [12]). This offset leads to cropping of the projections, see Fig. 4 a top row for full frames and Fig. 4 a bottom row for cropped frames (as indicated by the white frames). (An additional cropping is introduced by the offset between the Gaussian reference stack used for calibration and the Bessel stacks.) For tomographic reconstructions, this offset was corrected for by computing an offset vector field. For this, one Bessel beam stack was used as a reference. For each centroid of each bead in the segmented Bessel stack the offset vector to the same bead imaged in the three other segmented Bessel stacks was computed. Interpolating the offset vectors using linear regression for all three stacks results in the offset vector field:

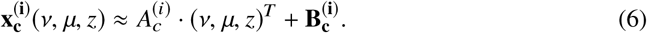

**Fig. 4.**
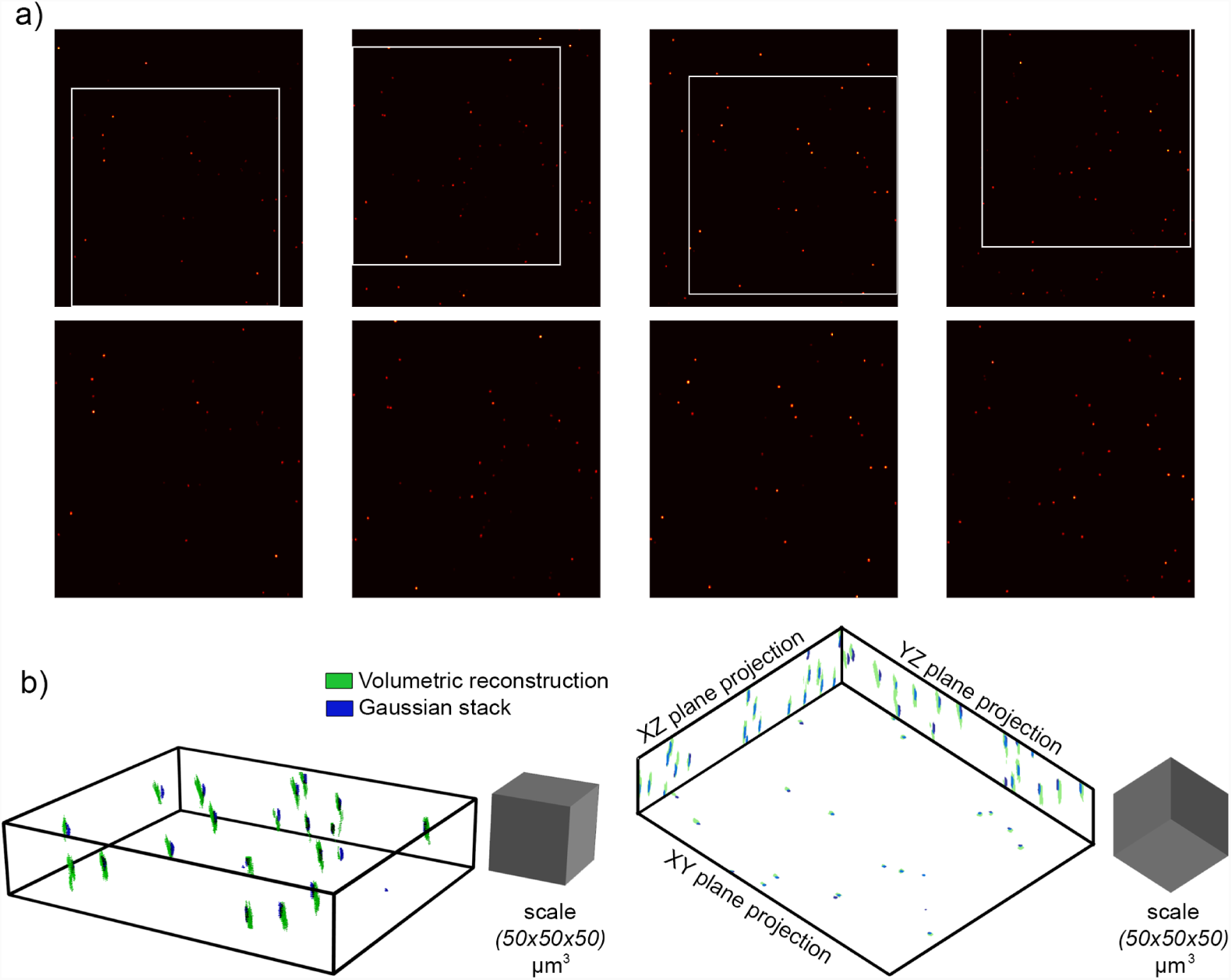
Tomographic reconstruction of 1 *μm* diameter beads. **a)** Top row: projections for each Bessel beam of a volumetric sample. The white frames indicate the overlap between the four Bessel beam projections and any bead within this area in visible in all projections. Bottom row: cropped projections corresponding to the overlapping area (white frames). **b)** Reconstruction of the beads (using only back projections, no convolutional networks) in green, compared to the volume recorded with the Gaussian beam (ground truth).

#### 2.2.3. Back Projection

Using the calibrated vector fields **x**_**c**_^(*i*)^, **ŝ** ^(*i*)^, and length *ℓ*^(*i*)^ for each projection *i*, the inverse radon transform, **ℜ**.^(−1)^***I***^(*i*)^, can be applied to any projection *I*^(*i*)^(*v, μ*), where the pixels of *I*^(*i*)^ are projected along the corresponding direction vector **ŝ** ^**(i)**^ and translated with its corresponding offset vector 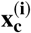. This results in a 3D image:

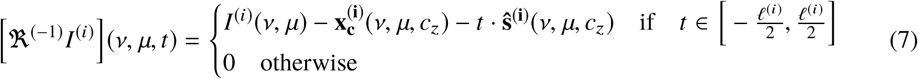

The parameter *t* needs to be discretized: *t = t*_*min*_, *(t*_*min*_ + *Z*_*res*_),…, *(t*_*max*_ *- Z*_*res*_), *t*_max_, where *z*_*res*_ gives the *z* resolution. We obtain an approximated reconstruction of the volume as the sum of the the inverse Radon transforms:

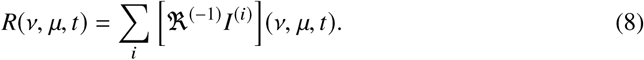

For small objects, such as the fluorescent beads in Fig. 4), we additionally filtered the back projections by setting the 3D image to zero if not all four inverse Radon transforms overlapped in a given position (*v, μ*, t). The recorded projections are shown in Fig. 4 a and the resulting reconstructed volume as well as a comparison with the Gaussian reference stack is shown in Fig. 4 b (no convolutional networks were applied for this reconstruction).

#### 2.2.4. Neural networks for distortion correction

For objects that are of similar size to the Bessel beam focal length, the back projected volume will extend over the entire volume due to the shallow projections angles. To recover axial resolution in this situation, neural networks were trained to correct for distortions and to recover axial resolution (see Fig. 1 b and c).

A three dimensional U-net (Fig. 5 a) was trained using simulated 3D samples consisting of ellipsoids with homogeneous constant density of different sizes (ranging from 1 *μm* to 40 *μm*) (Fig. 5 b). Using the Radon transform approach, four projections were generated from each simulated 3D sample using the vector fields measured during calibration. Shot noise was added in the four projections to simulate detection noise (Fig. 5 b, bottom row). From these four projections a (distorted) 3D volume was reconstructed using back projection (Fig. 5 b, left side). This reconstructed volume served as input to a 3D U-net (dimension 128*x*128*x*100 pixels) and the original simulated 3D sample (same dimension of 128*x*128*x*100) served as the output. The number of data samples generated was 1000.

**Fig. 5.**
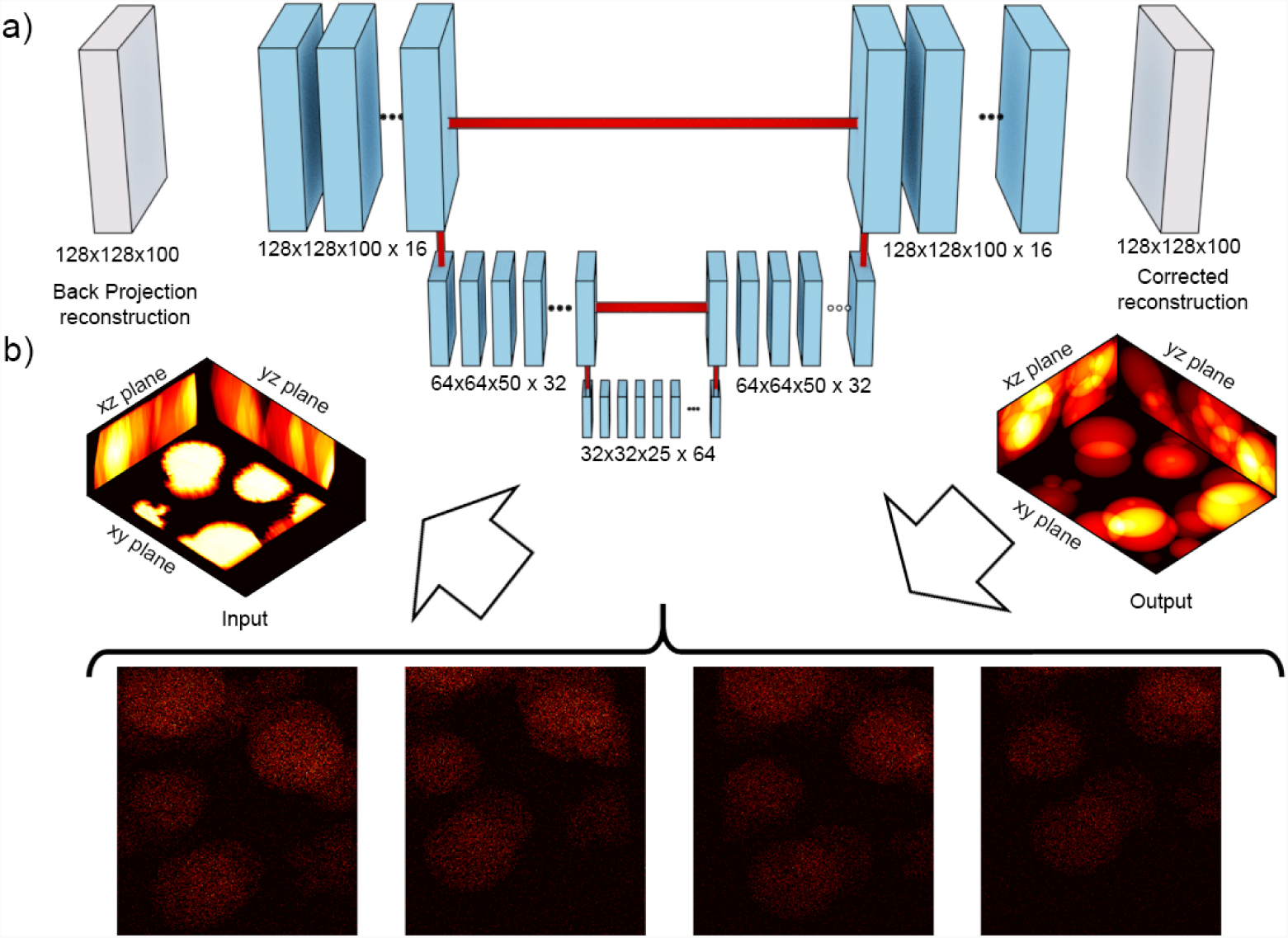
U-net architecture and simulated data. **a)** U-net: each step in the network is composed of 3D convolution, batch normalization and max pooling in 3D in the encoding part while the decoding part is composed of 3D convolution, batch normalization and up sampling in 3D. **b)** Simulated data used for training the network. The simulated 3D samples consist of ellipsoids of different sizes and serve as the output of the network. Four projections are generated from each simulated sample volume with addition of shot noise (bottom part). These four projections are produced using the calibration vector fields obtained experimentally. Applying back projection to these four projections yields a volume image. The black region in the reconstructed volume results from the cropping of the projections due to the offset between the different Bessel beam projections. This reconstructed volume serves as the input of the network, while the original volumes serves as the output.

Networks were built and trained using Python and *keras* with *tensor flow* as the backend [26]. The 3D U-net had 804.809 parameters and was composed of blocks formed in the first part by 3D convolutional layers, batch normalization and 3D max pooling. After the bottleneck we used a combination of 3D convolutional layers, batch normalization and 3D upsampling layers to deconvolve the 3D image and recover the corrected reconstruction (Fig. 5).

The model was trained on a desktop computer running Ubuntu with an Intel Xeon (R) CPU E5-1650 v4 @ 3.60GHz × 12 processor and a TITAN XP graphics card, that allowed to train the network for 100 epochs in about 5 hours. The loss during the training was selected to be the mean squared error between the original simulated image and the network output. We used the Adam optimizer to train the neural network. Results of applying this network on a pollen grain volume sample are shown in Fig. 6, where Fig. 6 a shows the four recorded projections and Fig. 6 b, top row, shows the reconstruction using back projection. Due to the - compared with the beam length - large size of the pollen grains, almost no axial resolution is recovered using back projection (Fig. 6 b, top row). However, axial information can be partially recovered with the U-net (Fig. 6 b, second row, compare to Fig. 6 b, last row for ground truth).

**Fig. 6.**
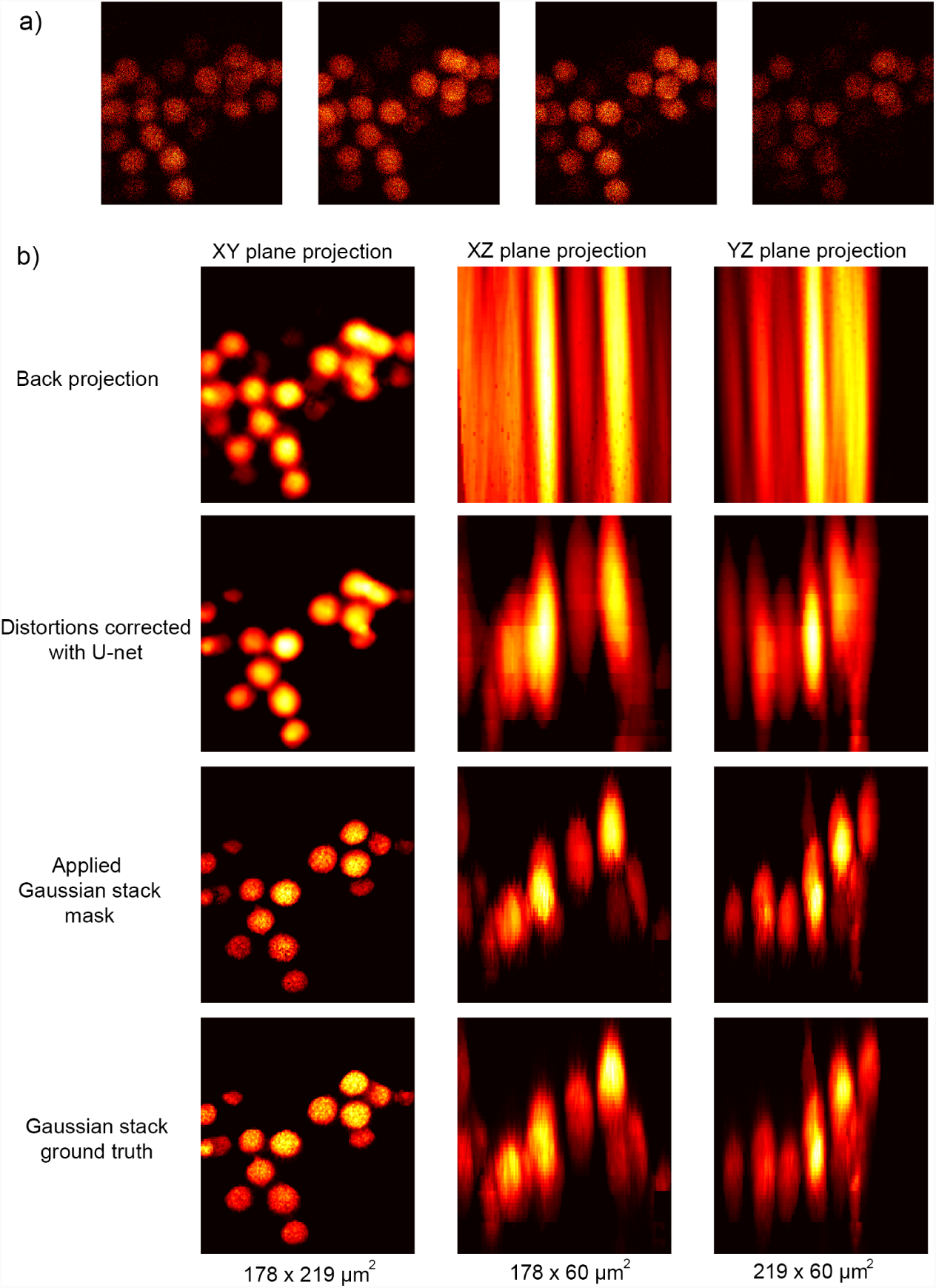
Tomography on pollen grain sample. **a)** Four projections recorded from a 3D sample of pollen grains. Only the overlapping region of the projections is considered for the reconstruction. **b)** Plane projections of the volumetric reconstructions. Top row: back projections show poor resolution in the *z* axis since the object approaches the beam length. The second row shows the reconstruction after applying the U-net. The third row additionally applies a mask recorded with the Gaussian beam (see text for details). The bottom row shows the reference Gaussian stack. The dimensions of the images are indicated for each image in the bottom row.

For sample which don’t change their structure during the observation time but only their intensity, for example neural activity in the brain, a Gaussian reference stack could first be recorded for obtaining structural information and Bessel beam tomography for subsequently monitoring activity at high temporal resolution [21]. We tested this approach by generating a structure-mask from the Gaussian stack by filtering it in the xy-plane with a Gaussian filter with 1 pixel standard deviation, thresholding it at the mean value, binarizing and downsampling it to 128 pixel resolution by averaging over four pixels. The resulting mask was multiplied with the volume reconstructed using combined back projection and U-net filtering (Fig. 6 b, second row) and resulted in the reconstruction shown in Fig. 6 b, third row. The accuracy of the reconstructed intensities depends on the quality of the tomographic reconstruction which in turn depends on the number of recorded projections and the reconstruction approach.

## 3. Discussion

We introduced a tomographic microscopy method for imaging volumes at frame scan rates. This was achieved using temporally multiplexed and tilted Bessel beams for recording independent projections at four different viewing angles with a single objective. A tomographic reconstruction approach suitable for scanning with tilted Bessel beams was developed combining Radon transforms with three dimensional convolutional networks (3D U-net). We recovered volume information in a single frame scan on fluorescent beads (Fig. 4) and pollen grains (Fig. 6). The frame rate depends on the resolution of the image and was 30 Hz for images with 512 pixel resolution (and increases with fewer pixels, for example 60 Hz for 256 pixel resolution).

Imaging using extended beams, lines or light sheets offers a promising approach for increasing the imaging rate beyond point scanning methods [1-4]. Different from approaches that detect fluorescence using a camera, Bessel beam tomography uses non-descanned PMT detection. This is an advantage for imaging in tissue, since non-descanned detection is not susceptible to scattering between different detection channels [3].

The depth of the field of view depends on the length of the Bessel beam and can be adjusted for the system of interest. For two-photon excitation, the limiting factor is the power that can be sent into the sample. The power could be optimized by using lasers with higher energy per pulse and lower repetition rate without exceeding the damage threshold [4]. Lower repetition rates would also reduce crosstalk between channels and would allow imaging with fluorophores with longer lifetimes. Bessel beam tomography experiments could alternatively also be performed using one- or three-photon excitation.

Different from previous approaches with tilted Bessel beams [9, 10, 12] we simultaneously recorded independent projections using temporal multiplexing. The advantage of simultaneous instead of sequential recordings [9, 10] lies in the increased frame rate and makes the measurement less sensitive to motion artefacts (which would interfere with the tomographic reconstruction) as typically observed in *in vivo* imaging. Increasing the number of projections to four is advantageous for volume reconstruction at shallow angles as illustrated in Fig. 7 and adding more projections is expected to only add a limited benefit for an isolated object. Different from another recent tomography approach [5], projections here were recorded along the optical axis instead of across the focal plane. While the axial projections at shallow angles come at the cost of lower resolutions it is compatible with a standard 2D scanning configuration and doesn’t require axial scanning.

**Fig. 7.**
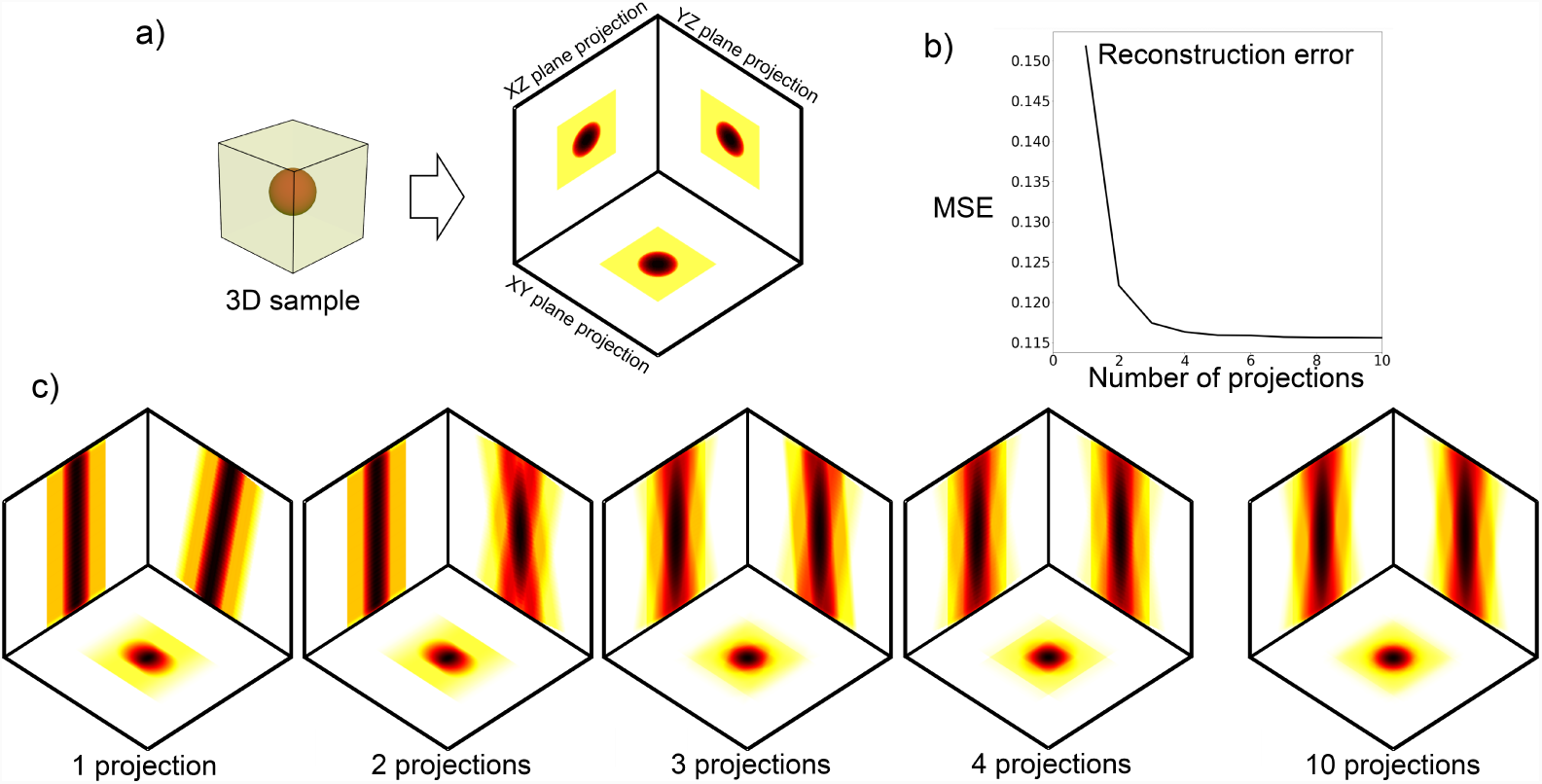
Simulated reconstruction error of 3D tomography depends on number of projections for the given isolated sample. **a)** A homogeneous cube that contains a homogeneous sphere with higher density and its projection in the *xy, xz* and *yz* planes (for visualization). **b)** Mean Square Error (MSE) between the simulated 3D sample and the reconstructed sample using back projection as a function of the number of recorded projections using a constant elevation angle of 170 degrees. **c)** Projections of the reconstructed sample considering 1,2,3,4 and 10 projections, respectively.

For volume reconstruction we used Radon inverse transforms combined with machine learning and adapted it for the presented Bessel beam scanning configuration. We modeled the Bessel beam as a constant line segment for computational simplicity, but models of the actual point spread function could be used for computing the inverse Radon transform. Approaches combining imaging physics and machine learning have been successfully applied in other tomography techniques for improving the reconstruction from sparse, shallow angle projections [13, 16-20]. Our networks were limited by GPU memory and therefore only low resolution images (128 pixel resolution instead of 512) were used. Training the models on higher resolution images will likely improve the reconstruction. More importantly, we used a very limited set of training samples which did not contain any high resolution features. Increasing the training dataset to include large numbers of more realistic samples and, if necessary, increasing the model complexity will allow to further improve the reconstruction and to recover higher resolution features.

Overall, Bessel beam tomography is suitable for fast imaging of sparse volumetric samples that can be accessed only with a single objective, a situation that is commonly encountered when imaging neural activity from cell bodies in the brain.

## Funding

Max Planck Society, caesar.

### Acknowledgments

We would like to thank Alex Turpin for initial experiments on Bessel beam generation, Georg Jaindl for Scanimage support, and Ivan Vishniakou for helpful discussions and comments on the manuscript.

## References

1. P. J. Keller and M. B. Ahrens, “Visualizing whole-brain activity and development at the single-cell level using light-sheet microscopy,” Neuron 85, 462–483 (2015).

2. J. A. Calarco and A. D. Samuel, “Imaging whole nervous systems: insights into behavior from worms to fish,” Nat. methods p. 1 (2018).

3. S. Weisenburger and A. Vaziri, “A guide to emerging technologies for large-scale and whole-brain optical imaging of neuronal activity,” Annu. review neuroscience (2018).

4. E. M. Hillman, V. Voleti, K. Patel, W. Li, H. Yu, C. Perez-Campos, S. E. Benezra, R. M. Bruno, and P. T. Galwaduge, “High-speed 3d imaging of cellular activity in the brain using axially-extended beams and light sheets,” Curr. Opinion neurobiology 50, 190–200 (2018).

5. A. Kazemipour, O. Novak, D. Flickinger, J. S. Marvin, J. King, P. Borden, S. Druckmann, K. Svoboda, L. L. Looger, and K. Podgorski, “Kilohertz frame-rate two-photon tomography,” bioRxiv p. 357269 (2018).

6. W. Amir, R. Carriles, E. E. Hoover, T. A. Planchon, C. G. Durfee, and J. A. Squier, “Simultaneous imaging of multiple focal planes using a two-photon scanning microscope,” Opt. letters 32, 1731–1733 (2007).

7. A. Cheng, J. T. Gonçalves, P. Golshani, K. Arisaka, and C. Portera-Cailliau, “Simultaneous two-photon calcium imaging at different depths with spatiotemporal multiplexing,” Nat. methods 8, 139 (2011).

8. G. Thériault, Y. De Koninck, and N. McCarthy, “Extended depth of field microscopy for rapid volumetric two-photon imaging,” Opt. express 21, 10095–10104 (2013).

9. G. Thériault, M. Cottet, A. Castonguay, N. McCarthy, and Y. De Koninck, “Extended two-photon microscopy in live samples with bessel beams: steadier focus, faster volume scans, and simpler stereoscopic imaging,” Front. Cellular neuroscience 8, 139 (2014).

10. Y. Yang, B. Yao, M. Lei, D. Dan, R. Li, M. Van Horn, X. Chen, Y. Li, and T. Ye, “Two-photon laser scanning stereomicroscopy for fast volumetric imaging,” PloS one 11, e0168885 (2016).

11. R. Lu, W. Sun, Y. Liang, A. Kerlin, J. Bierfeld, J. D. Seelig, D. E. Wilson, B. Scholl, B. Mohar, M. Tanimoto et al., “Video-rate volumetric functional imaging of the brain at synaptic resolution,” Nat. neuroscience 20, 620 (2017).

12. A. Song, A. S. Charles, S. A. Koay, J. L. Gauthier, S. Y. Thiberge, J. W. Pillow, and D. W. Tank, “Volumetric two-photon imaging of neurons using stereoscopy (vtwins),” Nat. methods 14, 420 (2017).

13. T. G. Feeman, “The mathematics of medical imaging,” Springer, (2010).

14. J. Sharpe, U. Ahlgren, P. Perry, B. Hill, A. Ross, J. Hecksher-Sørensen, R. Baldock, and D. Davidson, “Optical projection tomography as a tool for 3d microscopy and gene expression studies,” Science 296, 541–545 (2002).

15. M. Rieckher, U. J. Birk, H. Meyer, J. Ripoll, and N. Tavernarakis, “Microscopic optical projection tomography in vivo,” PLOS one 6, e18963 (2011).

16. X. F. Ma, M. Fukuhara, and T. Takeda, “Neural network ct image reconstruction method for small amount of projection data,” Nucl. Instruments Methods Phys. Res. Sect. A: Accel. Spectrometers, Detect. Assoc. Equip. 449, 366–377 (2000).

17. F. Thaler, C. Payer, and D. štern, “Volumetric reconstruction from a limited number of digitally reconstructed radiographs using cnns,” in Proceedings of the OAGM Workshop, (2018), pp. 13–19.

18. A. Goy, G. Roghoobur, S. Li, K. Arthur, A. I. Akinwande, and G. Barbastathis, “High-resolution limited-angle phase tomography of dense layered objects using deep neural networks,” arXiv preprint arXiv:1812.07380 (2018).

19. T. C. Nguyen, V. Bui, and G. Nehmetallah, “Computational optical tomography using 3-d deep convolutional neural networks,” Opt. Eng. 57, 043111 (2018).

20. D. Pelt, K. Batenburg, and J. Sethian, “Improving tomographic reconstruction from limited data using mixed-scale dense convolutional neural networks,” J. Imaging 4, 128 (2018).

21. R. Lu, M. Tanimoto, M. Koyama, and N. Ji, “50 hz volumetric functional imaging with continuously adjustable depth of focus,” Biomed. optics express 9, 1964–1976 (2018).

22. T. A. Pologruto, B. L. Sabatini, and K. Svoboda, “Scanimage: flexible software for operating laser scanning microscopes,” Biomed. engineering online 2, 13 (2003).

23. J. D. Seelig, M. E. Chiappe, G. K. Lott, A. Dutta, J. E. Osborne, M. B. Reiser, and V. Jayaraman, “Two-photon calcium imaging from head-fixed drosophila during optomotor walking behavior,” Nat. methods 7, 535 (2010).

24. O. Ronneberger, P. Fischer, and T. Brox, “U-net: Convolutional networks for biomedical image segmentation,” in International Conference on Medical image computing and computer-assisted intervention, (Springer, 2015), pp. 234–241.

25. Ö. Çiçek, A. Abdulkadir, S. S. Lienkamp, T. Brox, and O. Ronneberger, “3d u-net: learning dense volumetric segmentation from sparse annotation,” in International conference on medical image computing and computer-assisted intervention, (Springer, 2016), pp. 424–432.

26. F. Chollet et al., “Keras,” https://keras.io (2015).

